# The folate antagonist methotrexate diminishes replication of the coronavirus SARS-CoV-2 and enhances the antiviral efficacy of remdesivir in cell culture models

**DOI:** 10.1101/2020.07.18.210013

**Authors:** Kim M. Stegmann, Antje Dickmanns, Sabrina Gerber, Vella Nikolova, Luisa Klemke, Valentina Manzini, Denise Schlösser, Cathrin Bierwirth, Julia Freund, Maren Sitte, Raimond Lugert, Gabriela Salinas, Dirk Görlich, Bernd Wollnik, Uwe Groß, Matthias Dobbelstein

## Abstract

The search for successful therapies of infections with the coronavirus SARS-CoV-2 is ongoing. We tested inhibition of host cell nucleotide synthesis as a promising strategy to decrease the replication of SARS-CoV-2-RNA, thus diminishing the formation of virus progeny. Methotrexate (MTX) is an established drug for cancer therapy and to induce immunosuppression. The drug inhibits dihydrofolate reductase and other enzymes required for the synthesis of nucleotides. Strikingly, the replication of SARS-CoV-2 was inhibited by MTX in therapeutic concentrations around 1 μM, leading to more than 1000-fold reductions in virus progeny in Vero C1008 (Vero E6) as well as Calu-3 cells. Virus replication was more sensitive to equivalent concentrations of MTX than of the established antiviral agent remdesivir. MTX strongly diminished the synthesis of viral structural proteins and the amount of released virus RNA. Virus replication and protein synthesis were rescued by folinic acid (leucovorin) and also by inosine, indicating that purine depletion is the principal mechanism that allows MTX to reduce virus RNA synthesis. The combination of MTX with remdesivir led to synergistic impairment of virus replication, even at 300 nM MTX. The use of MTX in treating SARS-CoV-2 infections still awaits further evaluation regarding toxicity and efficacy in infected organisms, rather than cultured cells. Within the frame of these caveats, however, our results raise the perspective of a two-fold benefit from repurposing MTX for treating COVID-19. Firstly, its previously known ability to reduce aberrant inflammatory responses might dampen respiratory distress. In addition, its direct antiviral activity described here would limit the dissemination of the virus.

**SIGNIFICANCE:**

- MTX is one of the earliest cancer drugs to be developed, giving rise to seven decades of clinical experience. It is on the World Health Organization’s List of Essential Medicines, can be administered orally or parenterally, and its costs are at single digit € or $ amounts/day for standard treatment. In case of its successful further preclinical evaluation for treating SARS-CoV-2 infections, its repurposing to treat COVID-19 would thus be feasible, especially under low-resource conditions.
- Additional drugs exist to interfere with the synthesis of nucleotides, e.g. additional folate antagonists, inhibitors of GMP synthetase, or inhibitors of dihydroorotate dehydrogenase (DHODH). Such inhibitors have been approved as drugs for different purposes and might represent further therapeutic options against infections with SARS-CoV-2
- Remdesivir is currently the most established drug for treating COVID-19. Our results argue that MTX and remdesivir, even at moderate concentrations, can act in a synergistic fashion to repress virus replication to a considerably greater extent than either drug alone.
- COVID-19, in its severe forms, is characterized by pneumonia and acute respiratory distress syndrome, and additional organ involvements. These manifestations are not necessarily a direct consequence of virus replication and cytopathic effects, but rather a result of an uncontrolled inflammatory and immune response. Anti-inflammatory drugs such as glucocorticoids are thus being evaluated for treating COVID-19. However, this bears the risk of re-activating virus spread by suppressing a sufficient and specific immune response. In this situation, it is tempting to speculate that MTX might suppress both excessive inflammation as well as virus replication at the same time, thus limiting both the pathogenesis of pneumonia and also the spread of virus within a patient.

## INTRODUCTION

The SARS-CoV-2 pandemic has so far infected > 12 million people, with > 500,000 deaths ascribed to COVID-19 so far (July 2020). The fast spread of the virus, currently most notable in the Americas and south Asia, raises the need for readily available therapies, preferably through drug repurposing. In addition to viral proteins, cellular pathways represent attractive targets, with fewer opportunities for viruses to develop drug resistance.

The life cycle of SARS-CoV-2 comprises the replication of the viral RNA genome, one of the longest single strand RNA genomes among all viruses. Large amounts of RNA need to be synthesized, often subgenomic for translation at ribosomes, as well as full-length RNA genomes for incorporation into virus progeny. Thus, high amounts of ribonucleotides are required in infected cells for virus replication. This makes nucleotide biosynthesis an attractive target for drugs to interfere with the propagation of the virus in infected individuals.

One way of suppressing nucleotide biosynthesis consists in the use of folate antagonists. This class of drugs, upon cellular uptake, diminishes the regeneration of tetrahydrofolate (THF) from dihydrofolate (DHF), and the subsequent transfer of methyl groups required for the synthesis of purine nucleotides, as well as desoxythymidine monophosphate and S-adenosyl methionine. Especially the lack of purine nucleotides can be expected to diminish the synthesis of virus RNA. Folate antagonists have not been broadly evaluated as antivirals, with the exception of some flaviviruses (Beck et al., 2019; Fischer et al., 2013). In contrast, sulfonamides and trimethoprim are routinely used for treating bacterial infections, and their mechanism of action is to prevent the synthesis of folic acid in bacteria (Estrada et al., 2016).

Methotrexate (MTX), along with the closely related aminopterin, represents the most established antifolate to act on eukaryotic cells. They belong to the earliest cancer drugs whatsoever, since they were first characterized by Sidney Farber and colleagues in the forties of the last century (Farber et al., 1948), and have been in clinical use ever since. They were first applied at high doses to treat leukemia as well as solid malignancies. Later, however, they were found to suppress the inflammatory and immune response even at low doses (Cronstein and Aune, 2020). Until today, MTX is broadly used to treat rheumatoid arthritis and other autoimmune diseases, and it is also part of current standard anti-cancer regimens. It is bioavailable orally and parenterally, inexpensive, and, with seven decades of clinical use, one of the best-characterized drugs in the clinics. Its disadvantages are often a result of long-term treatment, which would not be necessary in the case of treating an acute virus infection. Of note, however, pregnancy is a contraindication for MTX, due to its negative impact on fast-proliferating embryonal cells (Weber-Schoendorfer et al., 2014). Moreover, high doses of MTX can cause severe mucositis (den Hoed et al., 2015). Thus, therapies with MTX do require close monitoring by trained personnel.

Here we show that MTX reduces SARS-CoV-2 replication up to 1000-fold, at concentrations that are clinically achievable, in Vero cells as well as Calu-3 cells that resemble bronchial epithelial cells. This effect of MTX can be largely ascribed to impaired purine synthesis. Furthermore, MTX synergizes with the antiviral purine analogue remdesivir to diminish virus replication. This raises the perspective that MTX might not only reduce inflammation in the context of COVID-19 but also display a direct antiviral impact.

## MATERIALS & METHODS

### Experimental Model And Methods

#### Cell culture

Vero E6 cells (Vero C1008) were kindly provided by C. Stahl-Hennig, German Primate Center Göttingen, Germany. Cells were maintained in Dulbecco’s modified Eagle’s medium (DMEM with GlutaMAX™, Gibco) supplemented with 10% fetal bovine serum (Merck), 50 units/ml penicillin, 50 μg/ml streptomycin (Gibco), and 10 μg/ml ciprofloxacin (Bayer) at 37°C in a humidified atmosphere with 5% CO_2_. The human lung epithelial cell line Calu-3 was purchased from ATCC (HTB-55) and maintained in Eagle’s Minimum Essential Medium (EMEM, Gibco) supplemented with 10% fetal bovine serum and penicillin/streptomycin. Cells were routinely tested and ensured to be negative for mycoplasma contamination.

#### Virus stock production

SARS-CoV-2 was isolated from a patient sample taken in March 2020 at Göttingen, Germany. Throat swab material was mixed with medium containing 2% fetal bovine serum and used to inoculate Vero E6 cells for five days. The supernatant was passaged twice more to expand the virus. This cell culture supernatant was used as virus stock in all experiments.

#### Treatments and SARS-CoV-2 infection

30,000 cells per well were seeded into 24-well-plates using medium containing 2% fetal bovine serum and incubated for 8 hours at 37 °C. Cells were treated with methotrexate (Selleckchem S1210), remdesivir (Selleckchem S8932), leucovorin (Selleckchem S1236) and inosine (Selleckchem S2442) at the concentrations indicated in the figure legends. When preparing stock solutions, methotrexate and remdesivir were dissolved in DMSO; leucovorin and inosine in water. After 24 hours, the cells were infected with virus stocks corresponding to 3*10^7^ RNA-copies of SARS-CoV-2 and incubated for 48-72 hours at 37 °C.

#### Quantitative RT-PCR for virus quantification

For RNA isolation, the SARS-CoV-2 containing cell culture supernatant was mixed (1:1 ratio) with the Lysis Binding Buffer from the Magnapure LC Kit # 03038505001 (Roche) containing > 3 M guanidine thiocyanate (GTC), and the viral RNA was isolated using Trizol LS. After adding chloroform, the RNA-containing aqueous phase was isolated, and RNA was precipitated using isopropanol. The RNA pellet was washed with ethanol, dried and resuspended in nuclease-free water. Quantitative RT-PCR was performed according to a previously established RT-PCR assay involving a TaqMan probe (Corman et al., 2020), to quantify virus yield. The following primers were purchased from Eurofins:

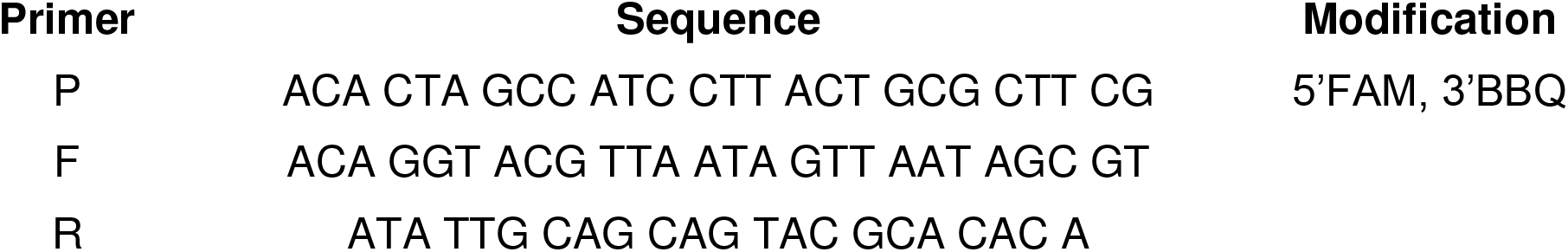

#### Immunofluorescence analyses

8,000 Vero E6 cells per well were seeded onto 8-well chamber slides (Nunc) and treated/infected as indicated. After 48 hours, the cells were fixed with 4% formaldehyde in PBS for 1 hour at room temperature, permeabilized with 0.5% Triton X-100 in PBS for 30 minutes and blocked in 10% FBS/PBS for 10 minutes. Primary antibodies were used to stain dsRNA (SCICONS J2 #10010200, 1:500) as well as the SARS-CoV-2 Spike (S; GeneTex #GTX 632604, 1:2000) and Nucleoprotein (N; Sino Biological #40143-R019, 1:5000) overnight. Secondary Alexa Fluor 488 donkey anti-mouse IgG and Alexa Fluor 546 donkey anti-rabbit IgG (Invitrogen, 1:500) antibodies were added together with 4′,6-diamidino-2-phenylindole (DAPI) for 1.5 hours at room temperature. All antibodies were diluted in blocking solution. Slides were mounted with Fluorescence Mounting Medium (DAKO) and fluorescence signals were detected by microscopy (Zeiss Axio Scope.A1).

#### ATP quantification by luminometry

30,000 Vero E6 cells per well were seeded into 24-well plates, treated with DMSO or MTX and infected with SARS-CoV-2 as indicated. After 48 hours, the cells were lysed with the Lysis Binding Buffer from the Magnapure LC Kit 03038505001 (Roche) containing > 3 M guanidine thiocyanate (GTC), and ATP levels were quantified by adding luciferase as well as luciferin to a 1 μl sample of the lysate, using 100 μl of the CellTiter-Glo^®^ Luminescent Cell Viability Assay solution (Promega), followed by luminometry using a Centro LB 960 luminometer (Berthold).

#### Immunoblot analysis

Cells were washed once in PBS and harvested in radioimmunoprecipitation assay (RIPA) lysis buffer (20 mM TRIS-HCl pH 7.5, 150 mM NaCl, 10 mM EDTA, 1% Triton-X 100, 1% deoxycholate salt, 0.1% SDS, 2 M urea) in the presence of protease inhibitors. Samples were briefly sonicated to disrupt DNA-protein complexes. The protein extracts were quantified using the Pierce BCA Protein assay kit (Thermo Fisher Scientific). Protein samples were boiled at 95 °C in Laemmli buffer for 5 min, and equal amounts were analyzed by sodium dodecyl sulfate polyacrylamide gel electrophoresis (SDS-PAGE). Subsequently, proteins were transferred onto a nitrocellulose membrane, blocked in 5% (w/v) non-fat milk in PBS containing 0.1% Tween-20 for 1 hour and incubated with primary antibodies at 4 °C overnight followed by incubation with peroxidase-conjugated secondary antibodies (donkey anti-rabbit or donkey anti-mouse IgG, Jackson Immunoresearch). The proteins were detected using either Super Signal West Femto Maximum Sensitivity Substrate (Thermo Fisher) or Immobilon Western Substrate (Millipore).

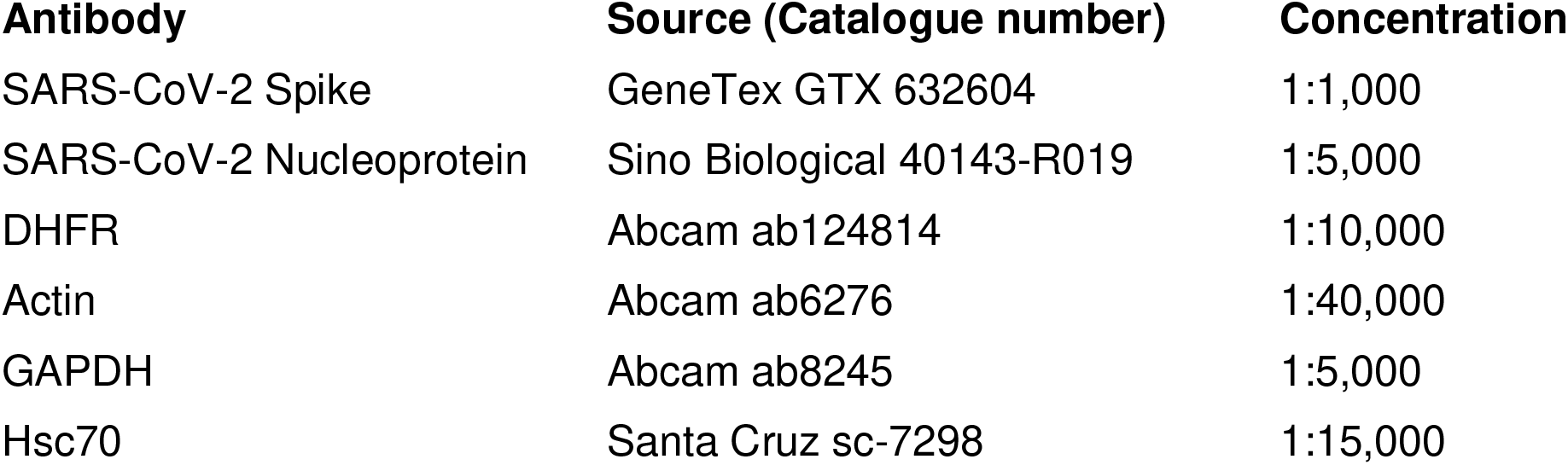

#### Cell counts (Celigo)

Vero E6 cells were seeded into 24-well-plates and treated/infected as indicated. After fixation of the cells and Hoechst staining to visualize nuclear DNA, the numbers of nuclei were counted using the Celígo^®^ S Imaging Cytometer (Nexcelom Bioscience).

#### RNA Sequencing and NGS data analysis

Quality and integrity of RNA were assessed using the Fragment Analyzer (Advanced Analytical) and the standard sensitivity RNA Analysis Kit (DNF-471). RNA-seq libraries were prepared using a non stranded RNA Seq, massively-parallel mRNA sequencing approach (Illumina; TruSeq RNA Library Preparation Kit v2, Set A; 48 samples, 12 indexes, Cat. N°RS-122-2001). Specifically, we first optimized the ligation step diluting the concentration of adaptors to increase ligation efficiency (>94%), and we reduced the number of PCR cycles (10 cycles) to avoid PCR duplication artifacts as well as primer dimers in the final library product. Libraries were prepared on an automated device (Beckman Coulter’s Biomek FXP workstation). For accurate quantitation of cDNA **l**ibraries, a fluorometry-based system, the QuantiFluor™–dsDNA System (Promega) was used. The size of final cDNA libraries was determined using the dsDNA 905 Reagent Kit (Fragment Analyzer from Advanced Bioanalytical) revealing sizes of 280 bp on average. Libraries were pooled and sequenced on the Illumina HiSeq 4000 analyzer (PE; 1 × 2×150 bp; 80 Mio reads/sample). Sequence images were transformed using the software BaseCaller (Illumina) to obtain BCL files, which were demultiplexed to FASTQ files using bcl2fastq v2.20.

Analysis of sequencing data was performed on the high-performance computing cluster provided by the Gesellschaft für wissenschaftliche Datenverarbeitung mbH Göttingen (GWDG) using the software Xshell® 6 for Home/School (NetSarang Computer Inc.). FASTQ files were subjected to quality control using FASTQC/0.11.4 (Babraham Bioinformatics) prior to trimming using FASTX/0.0.14 (Hannon Lab) according to following thresholds: first base to keep: 15, last base to keep: 140. Trimmed read pairs were mapped against the SARS-CoV-2 Wuhan-Hu-1 reference genome (accession number: NC_045512.2) (Wu et al., 2020) using BOWTIE2/2.3.4.1 (Langmead and Salzberg, 2012) with the --very-sensitive end-to-end setting. Resulting SAM files were processed using SAMTOOLS/1.9 (Li, 2011). In short, SAM files were converted into BAM files using samtools view, followed by merging of technical replicates using samtools merge. The resulting BAM files were sorted according to read names using samtools sort with the -n option and mate-scores were added to each read pair using samtools fixmate. The resulting BAM files were subsequently resorted according to genomic coordinates using samtools sort, PCR duplicates were removed using samtools markdup with default settings, and the corresponding index files were generated using samtools index. Sequencing depth was determined using samtools depth and the average read coverage per base was calculated taking the average sequencing depth for all positions across the reference genome. Genomic tracks of mapped reads were calculated using the bamCoverage function of DEEPTOOLS/3.0.1 (Ramirez et al., 2016) with the following settings: -e 100, --ignoreDuplicate, --smoothLength 60, --binSize 20, --samFlagInclude 64. SNPs present in the isolated virus were identified using bcftools mpileup with default settings, followed by bcftools call with -mv and -Ob settings of BCFTOOLS/1.9 (Samtools). Identified variants were filtered and converted into VCF files using bcftools view with the -i ‘%QUAL>=150’ setting. Genomic tracks as well as VCF files were visualized using the Integrative Genomics Viewer v2.3 (Robinson et al., 2017; Robinson et al., 2011; Thorvaldsdóttir et al., 2013).

### Quantification And Statistical Analysis

Statistical testing was performed using Graph Pad Prism 6 (RRID:SCR_002798). A two-sided unpaired Student’s t-test was calculated. Significance was assumed where p-values ≤ 0.05. Asterisks represent significance in the following way: ****, p ≤ 0.0001, ***, p ≤ 0.005; **, p ≤ 0.01; *, p ≤ 0.05.

## RESULTS

### Infection with SARS-CoV-2 produces >10^6^ copies of genomic virus RNA per cell

We isolated SARS-CoV-2 from a patient sample taken in March 2020 at Göttingen, Germany. Throat swab material was used to inoculate Vero C1008 cells (i.e. Vero E6 cells; shortly termed Vero cells from here on). The supernatant was passaged twice more to expand the virus, and cell supernatant was subjected to deep sequencing analysis. More than 80*10^6^ reads, 2*150bp each, were taken and aligned with the published genome sequence of the first Wuhan strain of SARS-CoV-2 (Wu et al., 2020). >97% of all reads corresponded to the virus. Comparing the Göttingen and Wuhan strains, only 8 nucleotide deviations were found (Fig. 1A). Most of these deviations were from C to U, arguing that they might have occurred through RNA deamination rather than misincorporation by the polymerase. Among the deviations was a nucleotide exchange leading to the mutation D614G in the Spike (S) protein that was recently reported to increase the number of S proteins displayed on the viral envelope (Li et al.; Zhang et al., 2020). Of note, however, no change in the coding region for the furin cleavage site of the S protein (Hoffmann et al., 2020; Liu et al., 2020; Ogando et al., 2020) was observed.

**FIGURE 1:**
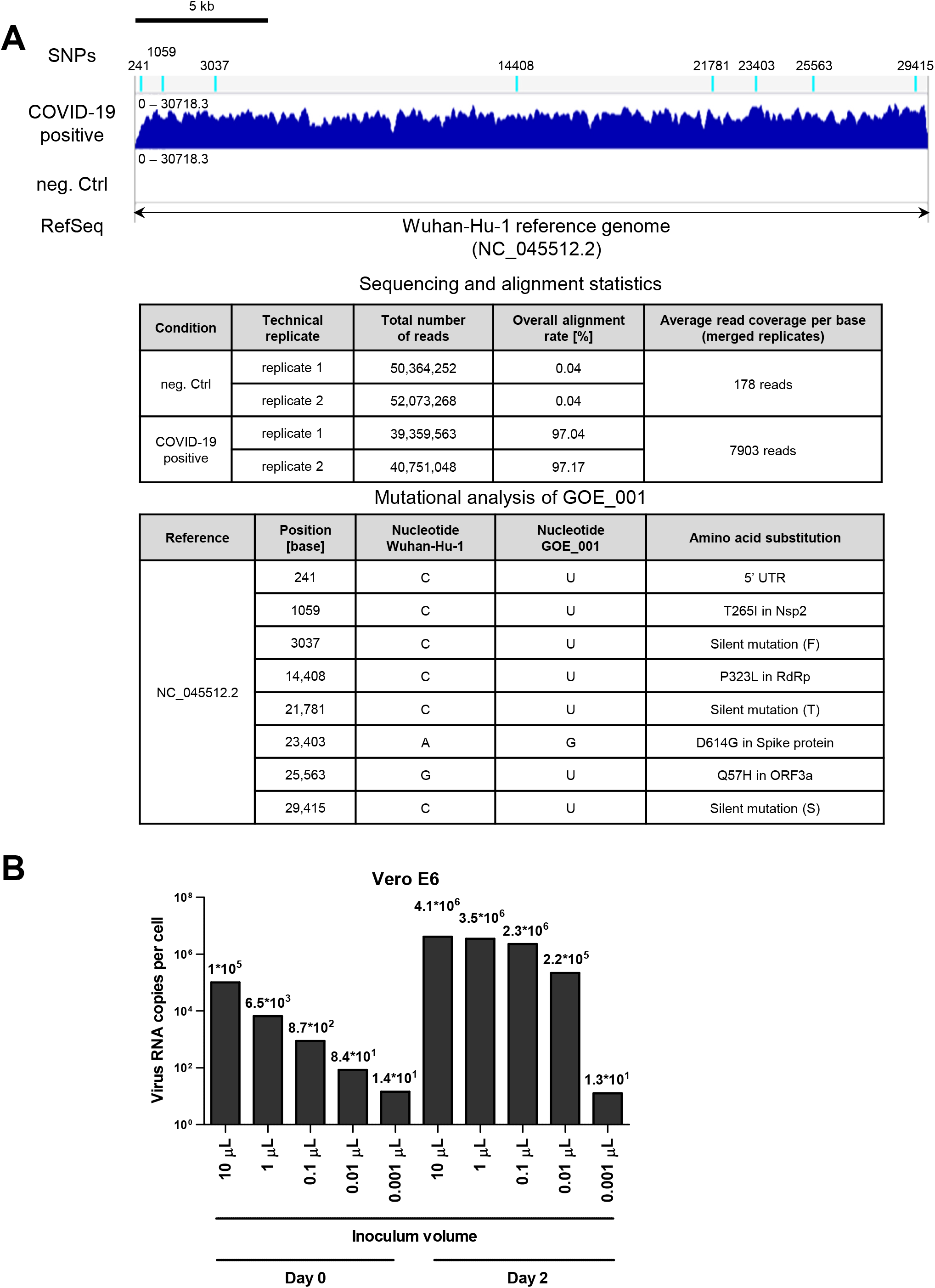
Characterization of a SARS-CoV-2 isolate. **(A)** Visualization of sequencing reads obtained from cell culture supernatants after infection with a sample of a COVID-19 positive patient (March 2020, Göttingen, Germany) and a non-infected cell monolayer (neg. Ctrl), mapped against the Wuhan-Hu-1 reference genome (NC_045512.2). Single nucleotide polymorphisms (SNPs) identified in the isolated SARS-CoV-2 strain (GOE_001) are indicated at the respective genomic positions as light blue bars on top of the read coverage. The tables underneath the genomic tracks describe the general sequencing and alignment statistics as well as the genetic alterations found. **(B)** Viral RNA copies found in the supernatant after inoculating 30,000 Vero cells with the indicated amounts of the primary virus stock. The amounts of RNA per infected cell were determined by RT-PCR in the day 0 inoculum and compared to the virus-derived RNA in the cell culture supernatant after two days of infection. Note that more than 106 RNA copies were synthesized and released to the media per infected cell.

Next, we detected and quantified the viral RNA released from infected cells. With decreasing virus inoculate, the yield of virus progeny after two days first remained at a maximum and then steeply decreased (Fig. 1B). The decrease in virus yield was more than linear compared to the decrease in inoculum. This could not solely be explained by the presence of defective virus particles. Rather, we propose that more than one virus particle might be required for the productive infection of a cell, leading to a more than proportional decrease in infectivity with lower virus concentrations. Quantitative RT-PCR with comparison to a standard (kind gift by the M. L. Schmidt, Robert Koch Institute, Berlin, Germany) revealed that up to 4*10^6^ copies of viral genomic RNA were released from each cell in the infected culture. Given the length of the viral genome of roughly 3*10^4^ nucleotides, this suggests that one infected cell needs to produce at least 3*4*10^10^ = 1,2*10^11^ nucleotides for virus RNA synthesis – and this does not include subgenomic virus RNA to encode virus proteins, nor negative strand RNA intermediates. Hence, virus production requires about 20 times more nucleotides than the replication of the cellular genome (6*10^9^ nucleotides in a diploid human cell). By this calculation, the demand of virus replication on nucleotide biosynthesis corresponds to more than three times of what is needed for the re-synthesis of all ribosomal RNAs, assuming 5*10^6^ ribosomes per cell (Lewin, 1994; Milo et al., 2010) and 7*10^3^ ribonucleotides per ribosome. Considering these numbers, we reasoned that nucleotide biosynthesis might represent a limiting factor for virus replication, and hence a suitable drug target.

### MTX strongly diminishes virus yield upon infection with SARS-CoV-2

As a strategy to interfere with nucleotide biosynthesis, and hence with virus replication, we treated Vero cells with MTX before and during infection with SARS-CoV-2. Strikingly, the treatment with MTX strongly antagonized the infection. MTX largely prevented the cytopathic effect (CPE) observed in infected cells, whereas up to 10 μM MTX did not produce obvious morphologic signs of cytotoxicity (Fig. 2A), in agreement with a previous preliminary report (Xing et al., 2020). The amount of virus RNA released into the media was reduced up to 1000-fold by MTX in Vero cells (Fig. 2B), and the effective concentration of MTX was around 1 μM, which was lower than the effective concentration of the established antiviral drug remdesivir, i.e. 3 μM (Fig. 2B). MTX diminished SARS-CoV-2 progeny even at 100 nM concentration when using Calu-3 cells (Fig. 2C), a cell line that can be used to model bronchial epithelia (Foster et al., 2000). Taken together, these results indicate that MTX is a potent inhibitor of SARS-CoV-2 replication.

**FIGURE 2:**
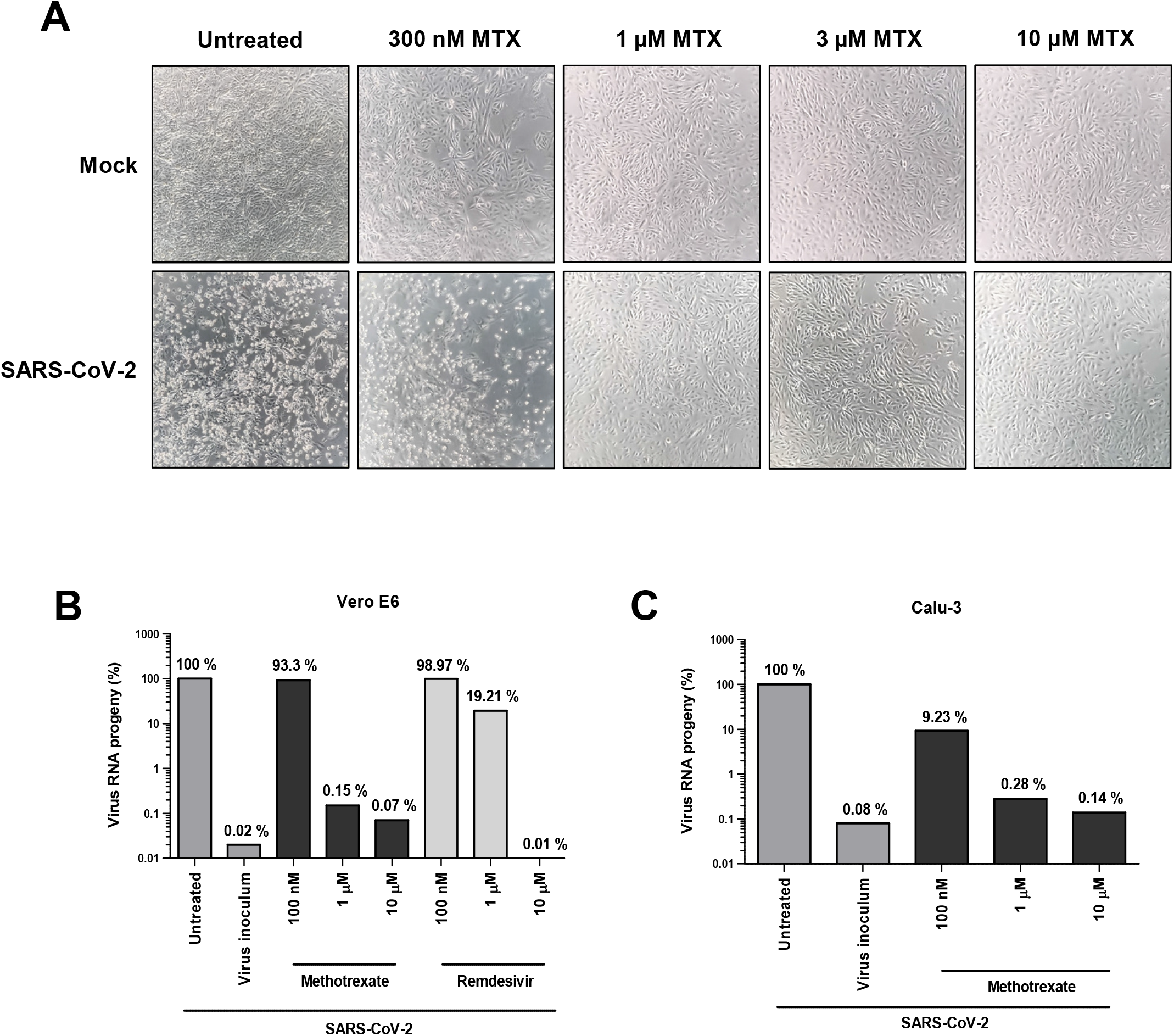
Impact of MTX on SARS-CoV-2 replication. **(A)** Diminished cytopathic effect (CPE) by MTX. 30.000 Vero cells were seeded, treated with MTX or the DMSO control, inoculated with the virus stock equivalent to 3*10^7^ copies of RNA and further incubated for 72 hrs. Cell morphology was assessed by phase contrast microscopy. Where indicated, the cells were treated with MTX at the indicated concentration for 24 hrs before and then throughout the time of infection. Note that the CPE was readily visible when comparing mock-infected and virus-infected cells, but only to a far lesser extent when the cells had been incubated with MTX. **(B)** Diminished virus progeny by MTX. Vero cells were treated with MTX and/or infected as above. At 48 hrs post infection (p.i.), RNA was prepared from the cell supernatant, followed by quantitative RT-PCR to detect virus RNA. The amount of RNA found upon infection without MTX treatment was defined as 100%, and the other RNA quantities were normalized accordingly. MTX was found capable of reducing virus RNA yield by more than 1000-fold. **(C)** Reduced virus yield by MTX in Calu-3 cells. Calu-3 cells were treated with MTX and/or infected as above. Again, MTX reduced the amount of virus RNA released to the supernatant by more than 1000-fold.

### In the presence of MTX, far lower amounts of double stranded RNA and virus proteins accumulate in SARS-CoV-2-infected cells

To obtain a first idea about the stage of the infectious cycle affected by MTX, we detected double stranded RNA (dsRNA; an intermediate of virus RNA replication) as well as the viral proteins, upon treating the cells with MTX and infecting them with SARS-CoV-2. The amount of viral S and N proteins, as well as the accumulation of dsRNA, were severely reduced by MTX (Fig. 3A, B). This argues that not just the release of virus particles is diminished by MTX, but also the earlier steps of the infectious cycle, i.e. the synthesis of RNA and virus proteins.

**FIGURE 3:**
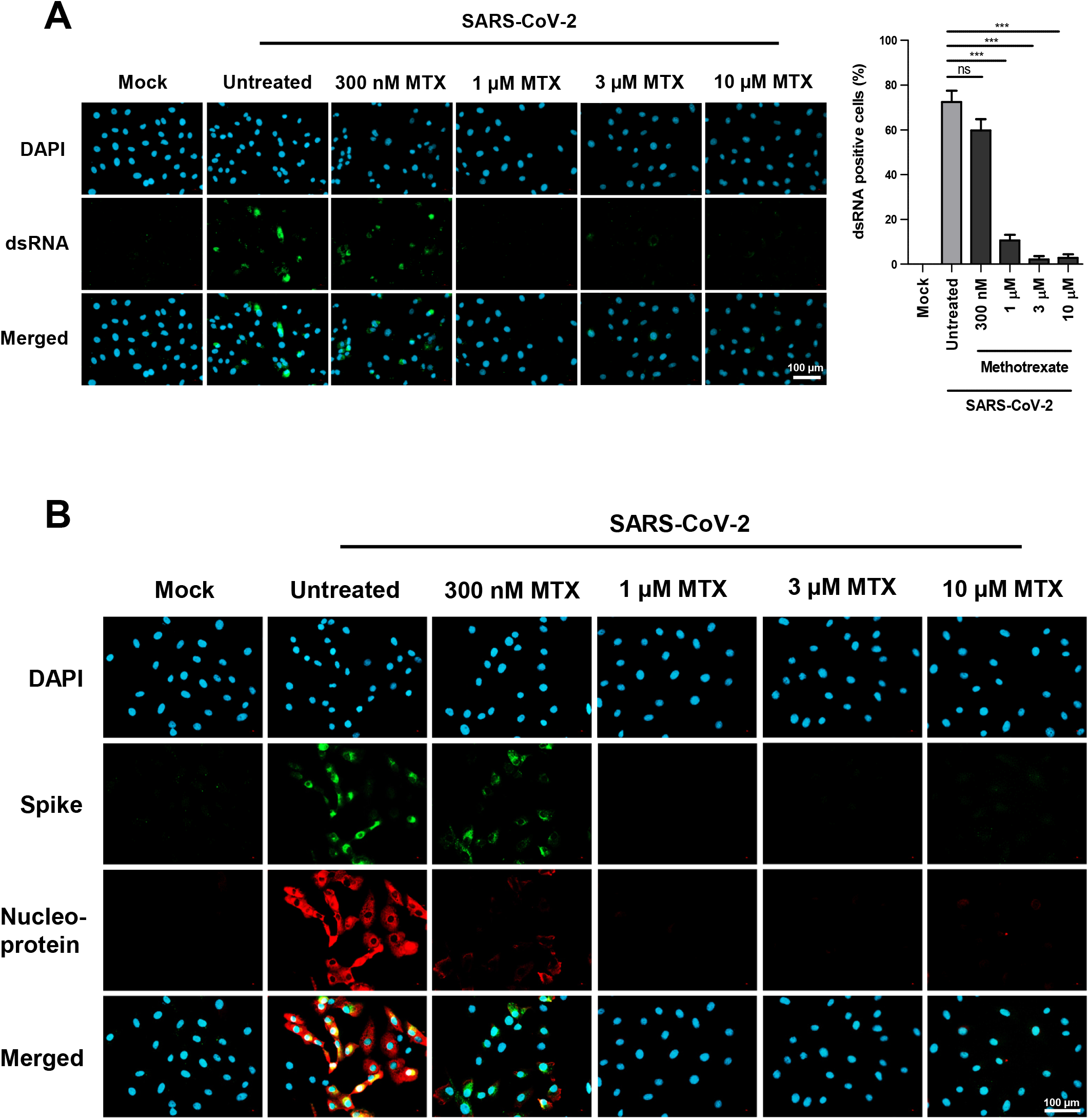
Reduction of dsRNA formation and viral protein synthesis by MTX. **(A)** Reduction of double stranded RNA (dsRNA) formation by MTX. Vero cells were treated with MTX as indicated and infected with SARS-CoV-2 for 48 hrs. DsRNA, a typical intermediate of virus RNA replication, was detected by immunofluorescence microscopy. **(B)** Reduction of viral protein synthesis by MTX. Cells were treated and infected as above. The SARS-CoV-2 Spike and Nucleoprotein were detected by immunofluorescence.

### MTX does not reduce the levels of its target dihydrofolate reductase in SARS-CoV-2-infected cells, but it strongly decreases the amount of cellular ATP

One of the major targets of MTX is dihydrofolate reductase (DHFR). To test if MTX can affect the levels of DHFR, we performed immunoblot analyses of Vero cells that were infected with SARS-CoV-2 and/or treated with MTX. We found infected cells to express the virus N and S proteins, whereas the amounts of these virus proteins were strongly diminished by MTX (Fig. 4A), as expected from the corresponding immunofluorescence analyses shown in Fig. 2B. In contrast, the levels of DHFR were not grossly affected by virus infection. However, MTX led to an increase of detectable DHFR, in mock-infected as well as virus-infected cells. Such an increase of DHFR upon MTX treatment has been described before and might be explained by increased translation and/or stability of DHFR in the presence of a folate antagonist (Hsieh et al., 2009). This observation suggests that SARS-CoV-2 infection does not overcome the adaptive increase in DHFR levels upon treatment of cells with MTX.

**FIGURE 4:**
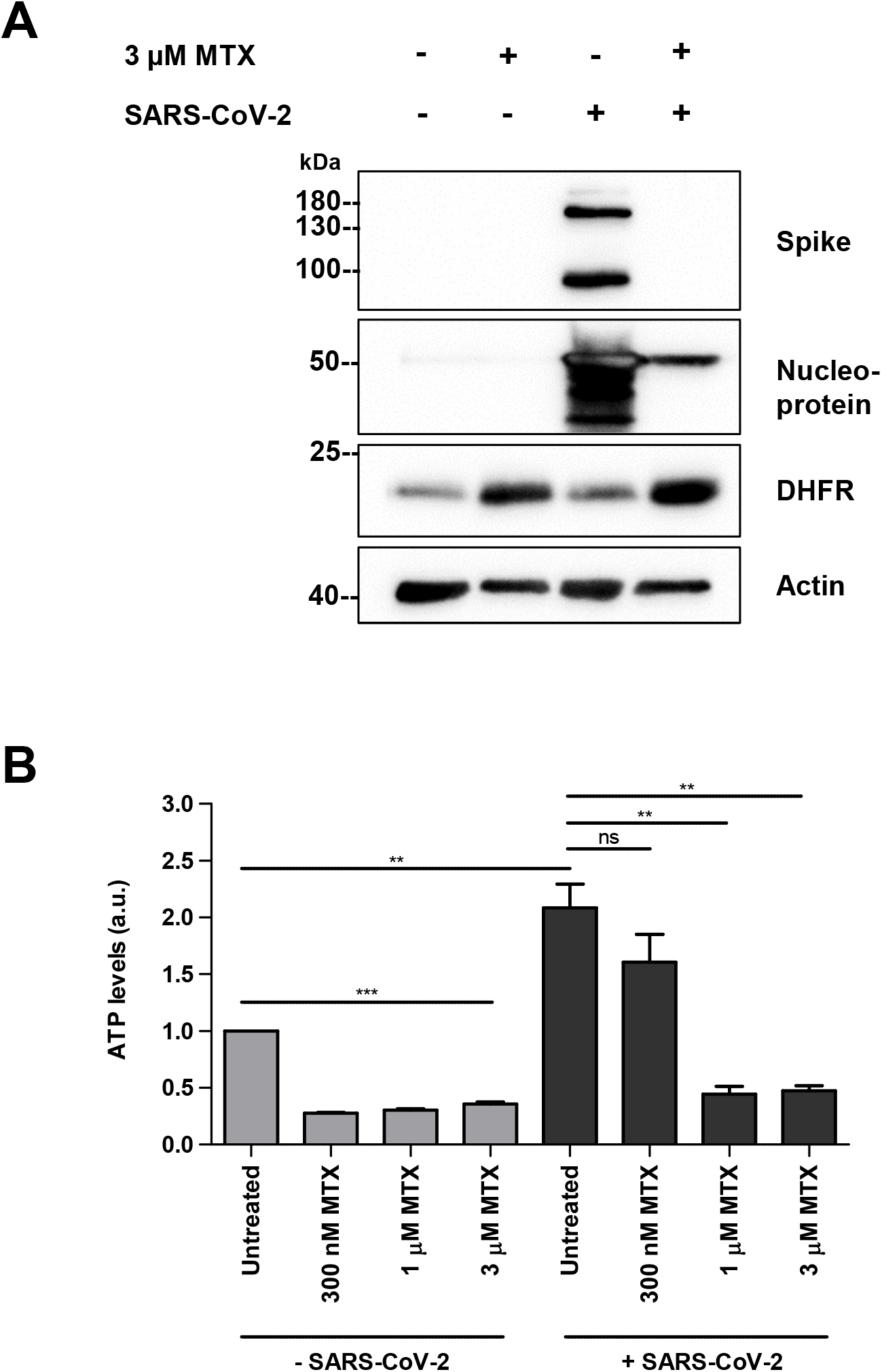
Levels of dihydrofolate reductase and ATP upon treatment with MTX and virus infection. **(A)** Increased levels of dihydrofolate reductease (DHFR) upon MTX treatment, regardless of virus infection, accompanied by decreased virus protein synthesis in the presence of MTX. Upon MTX treatment and/or infection of Vero cells with SARS-CoV-2 for 48 hrs, the viral N and S proteins as well as DHFR and actin (loading control) were detected by immunoblot analysis. Cells were treated with MTX as indicated and infected with SARS-CoV-2 for 48 hrs. **(B)** ATP levels upon MTX-treatment and SARS-CoV-2 infection. Vero cells were treated with MTX as indicated and infected with SARS-CoV-2 for 72 hrs. ATP levels were measured by the CellTiter-Glo^®^ assay. The extent of luminescence, reflecting relative ATP levels, was normalized to the number of cells found under the same conditions. Thus, the columns reflect the relative amount of ATP per cell in each case.

Next we asked whether purine synthesis might be diminished by MTX treatment in the context of infection. One key product of purine synthesis is represented by ATP. To determine ATP levels in MTX-treated and/or virus-infected cells, we rapidly lysed the cells in a chaotropic buffer, followed by dilution and addition of luciferin and luciferase to cell lysates, and luminometry. In this assay system, the levels of ATP are directly reflected by the extent of luminescence. Simultaneously, we determined the corresponding cell numbers by DAPI-staining of cell nuclei and automated quantification. When calculating the relative amount of ATP per cell, we found that MTX reduced ATP levels about three-to fivefold (Fig. 4B). Interestingly, ATP levels per cell were about two-fold higher in infected vs. non-infected cells, and reductions by MTX were only seen at higher concentrations of MTX. Nonetheless, MTX at 1 μM or higher concentration was capable of reducing ATP levels more than 5-fold in infected cells, suggesting that MTX impairs purine synthesis even upon infection with SARS-CoV-2.

### Folinic acid as well as purine nucleoside rescue virus replication in the presence of MTX

MTX interferes with a number of metabolic steps (Fig. 5A) after cellular uptake and polyglutamation (Alqarni and Zeidler, 2020; Bedoui et al., 2019; Cronstein and Aune, 2020). First and foremost, it inhibits dihydrofolate reductase (DHFR) (Nichol and Welch, 1950; Osborn et al., 1958). However, purine synthesis is inhibited by MTX even under conditions when the directly required folate metabolite, 10-formyl-THF, is largely preserved in its levels (Allegra et al., 1986). This argues that MTX diminishes purine synthesis, at least to a substantial extent, by direct inhibition of one of its metabolic steps. And indeed, polyglutamated MTX inhibits phosphoribosylaminoimidazolecarboxamide (AICAR) transformylase (ATIC) (Allegra et al., 1985b; Baggott et al., 1986).

**FIGURE 5:**
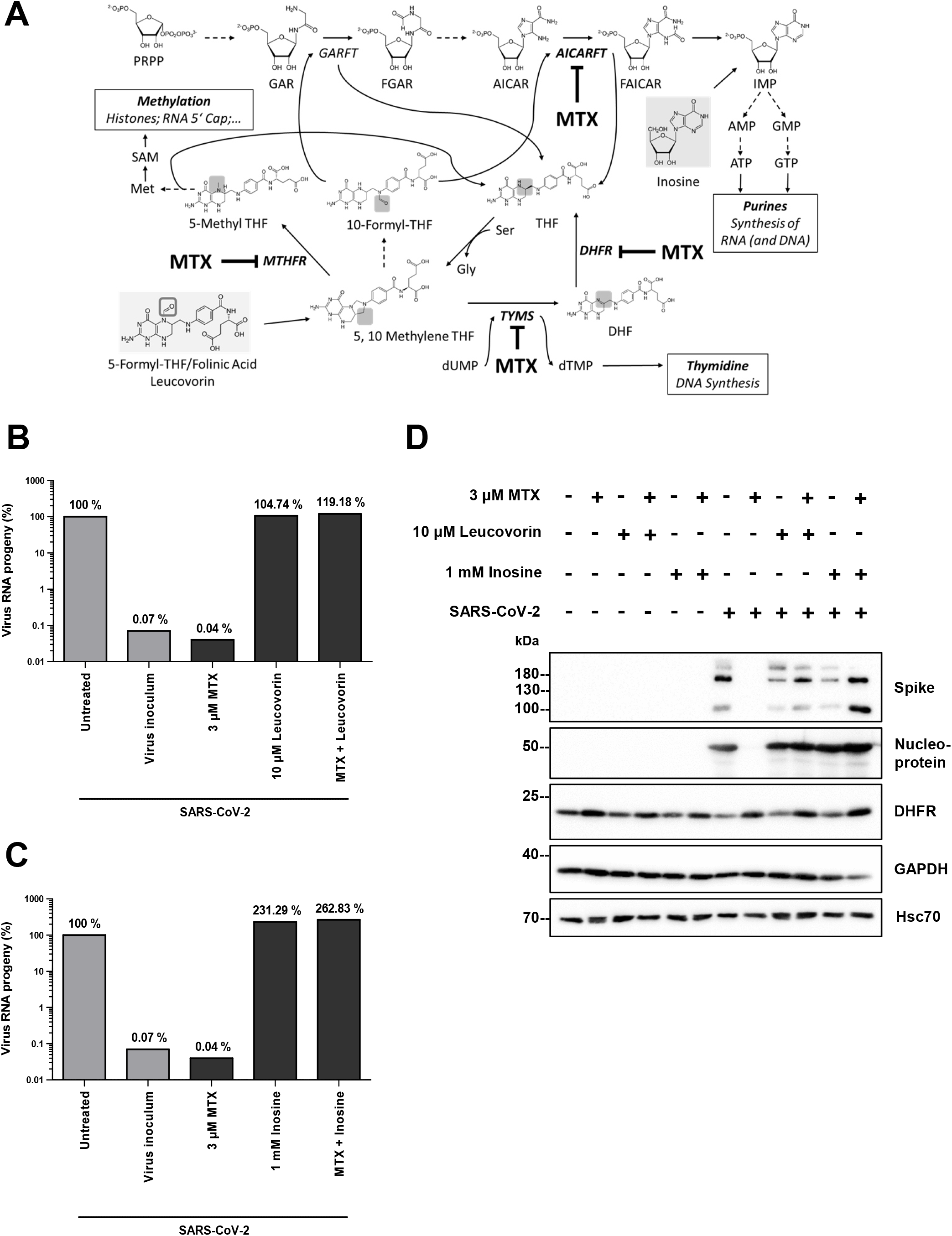
Rescue of virus replication by folinic acid and purine nucleoside upon MTX treatment. **(A)** Metabolic map indicating the major steps inhibited by MTX. *Enzymes* are indicated in *italics*. ***Enzymes directly inhibited by MTX*** are printed in ***bold and italics***. Plain arrows indicate direct reactions, dashed arrows indicate reactions with intermediates that are omitted for simplicity. Metabolites used for rescue experiments are shown with a grey background. <u>Folic acid metabolites</u>

- DHF, dihydrofolate
- THF, tetrahydrofolate <u>Folic acid metabolism, enzymes</u>

- ***DHFR***, dihydrofolate reductases
- ***MTHFR***, methylene tetrahydrofolate reductase
- ***TYMS***, thymidylate synthetase <u>Purine synthesis, enzymes</u>

- *GARFT*, glycinamide ribonucleotide formyltransferase
- ***AICARFT***, 5-aminoimidazole-4-carboxamide ribonucleotide formyltransferase, part of the dual function enzyme ATIC (comprising AICARFT and IMP cyclohydrolase) <u>Purine synthesis, metabolites</u>

- PRPP, phosphoribosyl pyrophosphate
- GAR, glycinamide ribonucleotide
- FGAR, formylglycinamide ribonucleotide
- AICAR, 5-aminoimidazole-4-carboxamide ribonucleotide
- FAICAR, 5-formamidoimidazole-4-carboxamide ribonucleotide <u>Nucleotides</u> are indicated by standard abbreviations, i.e. IMP (inosine monophosphate), AMP, GMP, ATP, GTP, dUMP, dTMP <u>Amino</u> acids are abbreviated as Ser, Gly, Met. **(B)** Rescue of virus replication in the presence of MTX by folinic acid (leucovorin). Vero cells were inoculated with virus as in Fig. 2, with or without MTX treatment 24 hrs before and during infection. In parallel or in addition, leucovorin was added to the cell culture media, at a concentration of 10 μM. **(C)** Restored virus replication by inosine, in the presence of MTX. The experiment was carried out as in (B), with the addition of 1 mM inosine instead of leucovorin. **(D)** Rescue of virus protein synthesis by leucovorin or inosine, in the presence of MTX. Upon treatment and infection of Vero cell as in (B) and (C), cell lysates were subjected to immunoblot analysis.

MTX, when polyglutamated, also inhibits thymidylate synthetase (TYMS) directly (Allegra et al., 1985a; Borsa and Whitmore, 1969), which, along with the depletion of 5,10 methylene THF, further diminishes the synthesis of deoxythymidine monophosphate (Chu et al., 1990). This then reduces the availability of deoxythymidine triphosphate, which is required for the synthesis of DNA but not RNA. Finally, MTX also acts as an inhibitor of methylene tetrahydrofolate-reductase (MTHFR) (Chabner et al., 1985). In addition to the shortage of the MTHFR substrate 5,10 methylene THF, this direct inhibition reduces the regeneration of methionine from homocysteine. Methionine is required to synthesize S-adenosylmethionine (SAM), a necessary substrate of methylations of proteins (e.g. histones) as well as the RNA 5’-cap. Given these diverse mechanisms of action by MTX, we performed rescue experiments to elucidate the mechanism(s) required to interfere with virus replication.

First, we added folinic acid (leucovorin) to the cells along with the MTX treatment. Remarkably, this fully rescued virus replication. Folinic acid is readily converted to 5,10 methylene THF, thus re-supplying the source of methylation required by all MTX-inhibited reactions (Chan and Cronstein, 2010). The rescue strongly suggests that the impact of MTX on virus replication is indeed due to its competition with endogenous folic acid and its metabolism (Fig. 5B).

Moreover, the addition of inosine restored virus replication in the presence of MTX (Fig. 5C). Both leucovorin and inosine were also capable of re-establishing the synthesis of virus N and S proteins (Fig. 5D). Inosine is rapidly converted to inosine monophosphate (Chan and Cronstein, 2010; Kozminski et al., 2020), which otherwise represents the end point of purine synthesis. This indicates that purine synthesis, rather than the supply of thymidine or SAM, is critical for virus replication. From this, the reduced supply of ribonucleotides appears as the dominating mechanism of action allowing MTX to act in an antiviral fashion.

### Remdesivir synergizes with MTX to antagonize SARS-CoV-2 replication

Remdesivir is currently the most advanced antiviral drug for clinical use against SARS-CoV-2 infection (Wang et al., 2020). It resembles a purine nucleoside and acts as an adenosine analogue to inhibit viral RNA polymerase. Since MTX diminishes the amounts of available ATP in SARS-CoV-2-infected cells (Fig. 4B), we reasoned that it might increase the incorporation of remdesivir and hence its antiviral activity. To test this, we first determined suboptimal concentrations of MTX and remdesivir to moderately diminish virus replication. Then, we treated Vero cells with the drugs at these concentrations, alone or in combination, followed by infection with SARS-CoV-2. And indeed, while each drug alone reduced the virus RNA in the cell supernatant (reflecting viral progeny) by roughly 50%, the combination of both drugs led to a reduction by more than 90% (Fig. 6A). Similarly, virus proteins were still detectable when the cells had been treated with single drugs at these concentrations, but the drug combination virtually extinguished the corresponding immunofluorescence signals (Fig. 6B). We conclude that MTX enhances the efficacy of remdesivir, leading to synergistic therapeutic efficacy.

**FIGURE 6:**
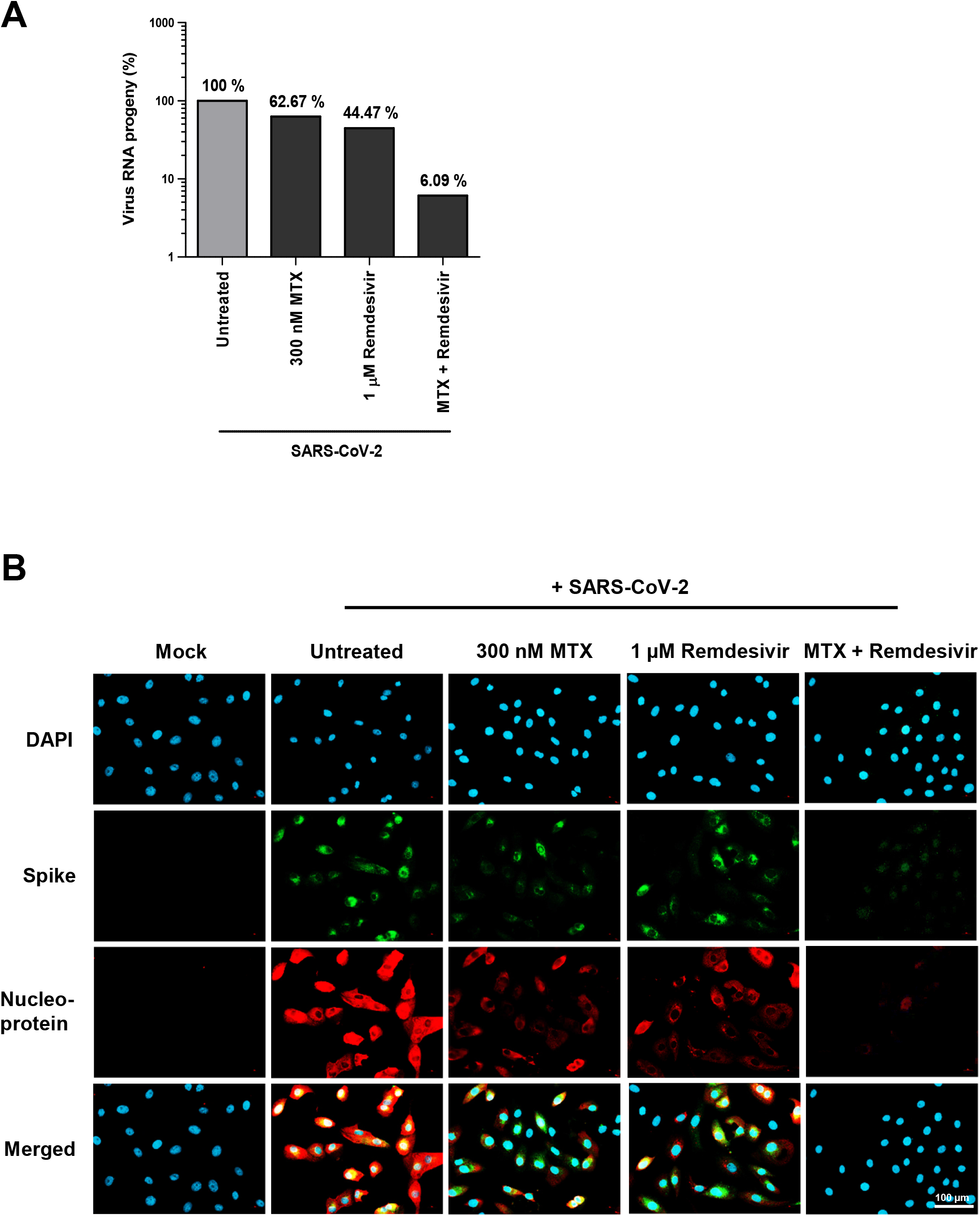
Synergism of MTX with remdesivir. **(A)** Remdesivir synergizes with MTX to antagonize SARS-CoV-2 replication. Vero cells were inoculated with virus as in Fig. 2, with or without MTX and remdesivir treatment 24 hrs before and during infection. **(B)** Stronger reduction of viral protein synthesis by a combination of MTX and remdesivir compared to single treatments, as determined 48 hrs p.i. by immunofluorescence analysis.

## DISCUSSION

Our results indicate that MTX efficiently inhibits the propagation of SARS-CoV-2, at least in the cell culture systems under study here. The inhibitory effect of MTX primarily relies on reduced purine synthesis. Moreover, MTX was combined with remdesivir to yield synergistic virus inhibition. Pending further testing in animals, these results are compatible with the idea of repurposing MTX as a drug for treating and/or preventing COVID-19, combining its known anti-inflammatory with the here-described antiviral effect.

The principle of limiting nucleotide synthesis to interfere with virus replication might be further expanded to treat COVID-19. To limit the synthesis of guanine nucleotides, the enzyme inosine-5′-monophosphate dehydrogenase (IMPDH) can be inhibited by mycophenolic acid, an established drug in the clinics (Allison and Eugui, 1996). Furthermore, pyrimidine synthesis can be reduced by inhibitors of orotate synthesis, targeting dihydroorotate dehydrogenase (DHODH). Besides teriflunomide, an approved immunomodulatory drug, novel DHODH inhibitors are in current clinical testing for treating cancer and leukemia (Madak et al., 2019) and might be re-purposed for treating SARS-CoV-2 infections as well, e.g. through a recently announced phase I trial (NCT04425252).

A reduction in available nucleotides may affect the synthesis of viral RNA not only through the kinetics of RNA polymerase. Given the strong impact of MTX on virus yield, with complete rescue by inosine, we speculate that low nucleotide concentrations may trigger regulatory signaling pathways for adapting the viral life cycle to such conditions. Such signaling pathways, as revealed by phosphoproteomics and metabolomics (Bojkova et al., 2020; Bouhaddou et al., 2020; Stukalov et al., 2020), might provide opportunities to further interfere with the replication of SARS-CoV-2.

In addition to purine synthesis, MTX also limits the transfer of C1 bodies (methyl groups) through MTHFR to regenerate S-adenosylmethionine (SAM), a major substrate for methyltransferases in the cell (Cronstein and Aune, 2020). Since SAM is required for the capping of the viral (as well as cellular) RNA (Viswanathan et al., 2020), this might further limit the production of infectious RNA. Indeed, the characterization of cellular metabolomics in response to SARS-CoV-2 infection, and the observed reduction in spermidine, at least suggested that SAM levels were reduced in infected cells (Gassen et al., 2020). However, based on the fact that the purine nucleoside inosine was capable of rescuing virus replication (Fig. 5C), we propose that the lack of purine synthesis represents the major mechanism of how MTX limits SARS-CoV-2 replication.

Using MTX against other viruses has been considered previously. As early as in 1957, it was reported that a folate antagonist was capable of protecting mice from infections with Lymphocytic Choriomeningitis Virus (LCMV; a member of the arenaviridae), while this was reverted by leucovorin (at that time still called “citrovorum factor”) (Haas et al., 1957). Moreover, MTX is capable of interfering with the replication of Zika virus, another positive strand RNA virus, and a member of the flaviviridae family of viruses (Beck et al., 2019). There, again, rescue with leucovorin and adenosine indicated reduced purine synthesis as the primary mechanism of action. However, at least to our knowledge, MTX has not been broadly used as an antiviral drug. We speculate that it might be usable against other viruses with large burst sizes and corresponding demand of nucleotides as well. Of note, other flaviviridae, i.e. Dengue and West Nile Virus, were also found susceptible to treatment with MTX, whereas Sindbis and Vesicular Stomatitis Virus were not (Fischer et al., 2013). It thus remains to be determined whether MTX might also interfere with the replication of other RNA as well as DNA viruses.

With seven decades of clinical experience, MTX is one of the most established drugs whatsoever, available at low costs, making its potential use for treating COVID-19 relatively straightforward. Having said this, a number of caveats still apply. Firstly, the dose of MTX that would be needed for antiviral treatment is probably higher than the typical dose taken by patients for the purpose of long-term anti-inflammation treatment. Plasma concentrations in patients treated for rheumatoid arthritis, typically taking 7.5 mg/week for months or even years, are only around 25 nM (Hornung et al., 2008), which is unlikely to interfere with virus replication. On the other hand, when applying high-dose therapies, up to 1000 μM plasma concentrations are achievable (Comandone et al., 2005), i.e. 1000-fold more than what would be required for the antiviral effect in our study. Under such circumstances, a limiting toxicity would consist in mucositis, which is often observed when treating leukemia with high-dose MTX in a pediatric setting (den Hoed et al., 2015).

Antiviral activities of MTX would presumably require an intermediate shortterm dose for a week or two. The antiviral concentration of MTX (> 1 μM) is achieved, for instance, by single intramuscular application of 50 mg MTX, which is a common procedure to induce abortions at early stages of pregnancy (Creinin and Vittinghoff, 1994). In this setting, tolerable side effects were reported, e.g. transient nausea, diarrhea, and headache (Creinin and Krohn, 1997). This comparison, on the other hand, makes it very clear that antiviral doses of MTX are contraindicated in pregnant women.

A clinical study using nanoparticle-coupled MTX for treating COVID-19 was registered at clinicaltrials.gov, NCT04352465. Moreover, clinical studies using intermediate-to-high dose therapy with MTX were recently proposed for COVID-19, purely based on the rationale of immunosuppression by MTX (Frohman et al., 2020a; Frohman et al., 2020b). Our findings that MTX also interferes with virus replication further argues in favor of such clinical studies.

As with other antiviral drugs such as remdesivir, the timing of the therapeutic intervention appears critical. At stages where the COVID-19 symptoms are fully established, virus propagation may already have surpassed its peak. The disease is then mostly caused by excessive inflammatory responses in the lung, as well as intravasal blood coagulation. This argues in favor of early treatment with drugs that interfere with virus replication, to avoid spreading of the infection in the first place.

On the other hand, MTX may benefit COVID-19 patients not only through its direct impact on virus replication. Rather, MTX has long been known to suppress the immune response by diminishing the proliferation of B and T cells, which may reduce the pathogenesis of immune-driven lung pathology. While immune suppression would otherwise bear the risk of further virus propagation, our results strongly suggest a direct impact of MTX on viral RNA synthesis. This might allow two benefits at once, i.e. lower virus spread and lower immune reaction. Finally, MTX not only reduces the adaptive immune response, but also the inflammatory reaction, e.g. through extracellular release of adenosine (Cronstein et al., 1991) as well as by diminishing the JAK-STAT signaling pathway (Thomas et al., 2015). Indeed, the targeted JAK-STAT inhibitor baricitinib was reported to be beneficial for COVID-19 patients (Richardson et al., 2020), and combining baricitinib with MTX was suggested recently (Seif et al., 2020); furthermore, the reduction of the inflammatory response may also explain the benefits of corticosteroids in the management of COVID-19 (Li et al., 2020).

So far, the use of MTX in patients with rheumatoid arthritis was at least reported to be of no detrimental effect when these patients had COVID-19 (Sanchez-Piedra et al., 2020). However, none of these studies was aiming at a direct antiviral effect of the drugs. According to the results presented here, MTX would have the additional advantage of interfering with virus replication itself.

The clinical efficacy of MTX against SARS-CoV-2 replication might be further enhanced through the combination with antivirals such as remdesivir. We propose that this combination might work in a more than additive fashion, but by a mechanistic synergism. Specifically, the reduction in available purine nucleotides might enhance the likelihood of incorporating remdesivir or similar antiviral drugs and their active metabolites into the nascent viral RNA. In any case, treatment with remdesivir andMTX might not only be tolerable but serve to potentiate antiviral efficacy.

In summary, albeit solely based on cell culture experiments so far, our study raises the possibility of using MTX, alone or in combination with remdesivir, to limit the replication of SARS-CoV-2 in patients, with the possible additional benefit of its immunosuppressive and anti-inflammatory effects to reduce the pathogenesis of COVID-19. Pending further preclinical evaluation, repurposing the established drug MTX for treating COVID-19 might be beneficial.

## ACKNOWLEDGEMENTS

We thank Stefan Pöhlmann, German Primate Center Göttingen, for helpful advice, and Marie Luise Schmidt, Institute of Virology, Charité Berlin, for SARS-CoV-2-derived RNA. Moreover, we thank Carsten Lüder and Melanie Eisele as well as the members of the Institute of Medical Microbiology for their continuous help and support in working at biosafety level 3. KMS and LK were members of the Göttingen Graduate School GGNB during this work. VN and VM were members of the IMPRS/MSc./PhD program Molecular Biology.

## AUTHOR CONTRIBUTIONS

MD conceived the project. MD, KMS and AD designed experiments. AD and KS performed most experiments. VN, LK, VM, DS, CB and JF established and/or performed virus quantification. MS, GS, BW and SG performed sequence analyses. RL, DG and UG provided guidance in virus isolation and quantification. MD wrote the manuscript. All authors read and approved the manuscript.

